# AutoGenome V2: New Multimodal Approach Developed for Multi-Omics Research

**DOI:** 10.1101/2020.04.02.021345

**Authors:** Chi Xu, Denghui Liu, Lei Zhang, Zhimeng Xu, Wenjun He, Deyong Wang, Mingyue Zheng, Nan Qiao

## Abstract

Deep learning is very promising in solving problems in omics research, such as genomics, epigenomics, proteomics, and metabolics. The design of neural network architecture is very important in modeling omics data against different scientific problems. Residual fully-connected neural network (RFCN) was proposed to provide better neural network architectures for modeling omics data. The next challenge for omics research is how to integrate informations from different omics data using deep learning, so that information from different molecular system levels could be combined to predict the target. In this paper, we present a novel multimodal approach that could efficiently integrate information from different omics data and achieve better accuracy than previous approaches. We evaluate our method in four different tasks: drug repositioning, target gene prediction, breast cancer subtyping and cancer type prediction, and all the four tasks achieved state of art performances. The multimodal approach is implemented in AutoGenome V2 and is also powered with all the previous AutoML convenience to facilitate biomedical researchers.

## Introduction

With the development of sequencing technologies, researchers have extended the large scale whole genome profiling experiments from genomics to epigenetics, proteomics and metabolics. Obtain whole omics profiling data from a single sample/individual are more and more popular in biomedical researches^1^, it helps researchers to extract evidences from different molecular system levels, to explore and understand the underlying biological mechanisms. For example, in cancer research, researchers need to confirm evidences from cancer cell gene expression, gene mutations, gene copy number variations and gene methylations to form a proper hypothesis, tools like oncoplot is developed to help visualize and analyze the multi-omics data^2^.

Deep learning is very popular in genomics research recently^3^. Previous work have proved the advantage of Deep Neural Network (DNN) against traditional machine learning methods, such as support vector machine (SVM), logistic regression and Xgboost^4–6^, in single-omics area. How to apply deep learning in multi-omics research, is a new challenging area for researchers.

The simplest way to integrate multi-omics data is to concatenate all the omics data as the input to the DNN. For example, DeepSynergy^7^ concatenate fingerprints of molecular structures of chemicals and omics data of cancer cell lines directly as input for the Multi-layer perception neuron network (MLP). The problem is that the data distributions in different omics data vary a lot, some omics data even have different data type, which makes the DNN difficult to fit a good model.

The advanced approach is to use sub-network for each omics data, and concatenate the output of the sub-networks to predict the targets. For example, MOLI^8^ uses three MLP sub-networks for the three omics data (GDSC^9^ cancer cell line gene expression, mutation and CNV), the three sub-networks are then concatenates together to predict the targets. PaccMann^5^ extended this approach by using three different sub-network to model molecular structure, omics data and gene interaction network separately. The limitation with this approach is that it cannot fine tuning or optimize the sub-networks independently.

The more sophisticated approach is to train DNN separately for each omics data, and then concatenate the embedding layers together to make the final predictions, for example, DeepDR^6^ train autoencoder (AE) for gene expression and gene mutation separately, the latent space are then concatenated and forwarded with MLP to make the final prediction.

The proposal of residual fully-connected neural network (RFCN) implemented in AutoGenome^10^ have shown us a good framework to model the single-omics data, we want to go further to extend it to multi-omics research, by considering a. the advantage of RFCN neural network architectures; b. the AutoML features from AutoGenome; c. novel multi-model approaches to integrate different multi-omics data. In this paper, AutoGenome is further developed for this purpose, users specify the location of the input omics data and the learning targets, AutoML is then used to train the DNN model automatically, after an optimal model is trained, model interpretation module will be used to explain the influence of each gene to the learning targets. AutoGenome supports both regression and classification tasks, it doesn’t require the users to master tensorflow^11^ or pytorch^12^ to start with.

We evaluate the performances of AutoGenome in four different multi-omics biomedical tasks: a. drug response prediction, b. gene dependency prediction, c. breast cancer subtype prediction and d. pan-cancer patient stratification, and AutoGenome outperformed all the existing methods. The results show that AutoGenome could efficiently integrate large scale multi-omics data and generate explainable AI models. We envision AutoGenome to become a popular tool in multi-omics research.

## Results

### Overview of AutoGenome

Researchers from hospitals, pharmaceutical companies and academic institutes usually use patient tissues, animal models and cell lines in their research to study biomedical problems. To uncovering the molecular level mechanisms, high throughput sequencing technologies are then used to profiling multiple types of omics data, such as gene expression, gene mutation, copy number variation (CNV), DNA methylation, microRNA and histone modification (Figure 1). By analyzing the multi-omics data, researcher could formulate new hypothesis or create mathematical models for forecasting, such as drug sensitivity prediction, gene dependency prediction, patient stratifications, and so on. Multi-omics data represent molecular phenotypes at different molecular systems, each omics data have different distributions, which makes it very challenging to model with.

**Figure 1.**
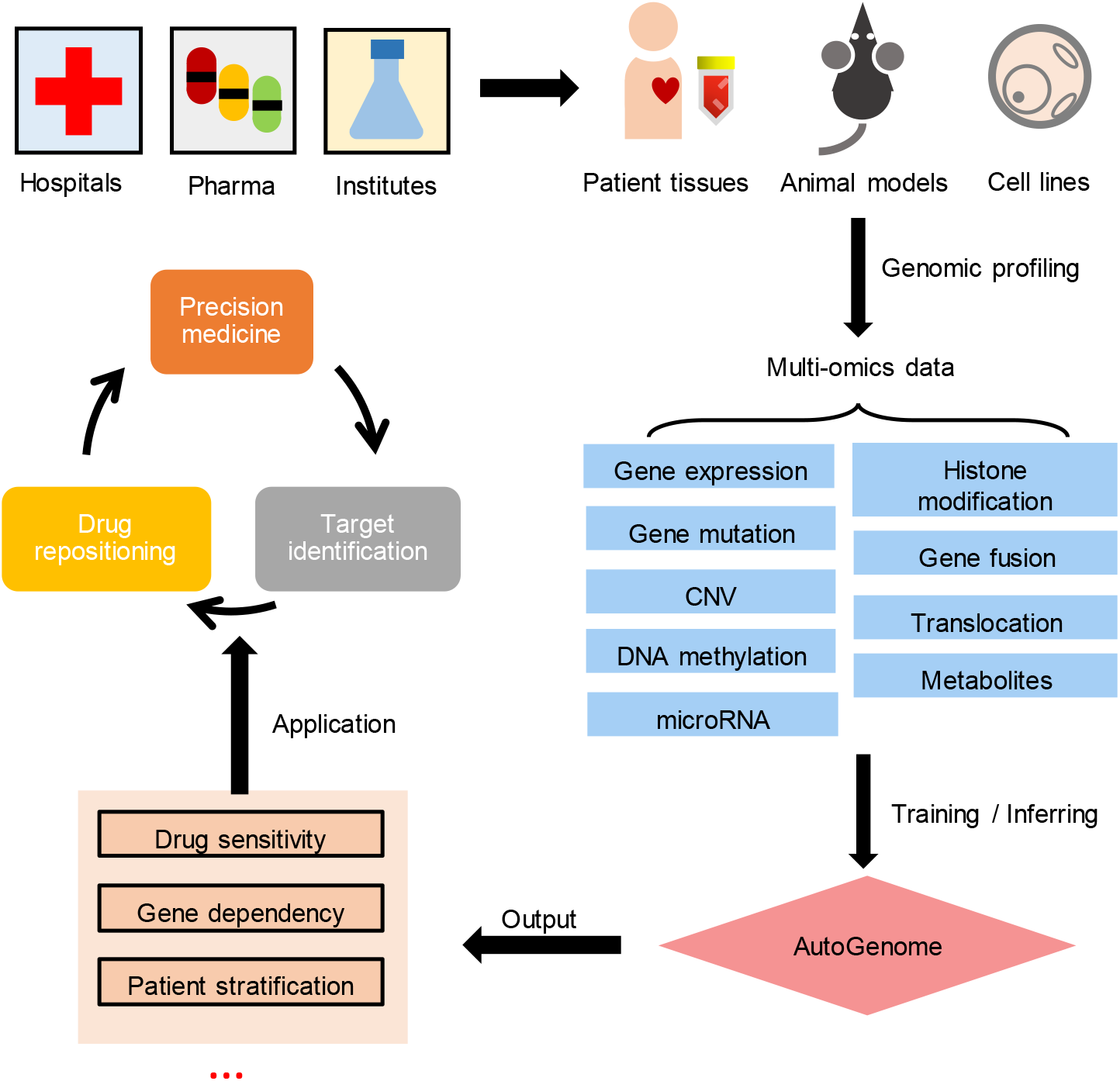
Integrative analysis of multi-omics data from multiple-origin samples and building AI models for biomedicine research.

AutoGenome provide a convenient framework to help researchers to build the best multi-model deep neuron network for their research. There are four steps in the pipeline (Figure 2). **Step 1**: Prepare DNN input, different omics data are prepared into normalized 2-dimensional matrixes with samples and genes. **Step 2**: AutoML is used to search for the best DNN model for each single-omics data against the learning target, the neuron network architectures used in AutoML search space including MLP, RFCN-ResNet, RFCN-DenseNet and Random-wired RFCN^10^ (Methods). **Step 3**: The last layers from the single-omics models are concatenate together as the input for the final multi-omics DNN model. **Step 4**: AutoML is used again to train the final multi-omics DNN model against the learning target.

**Figure 2.**
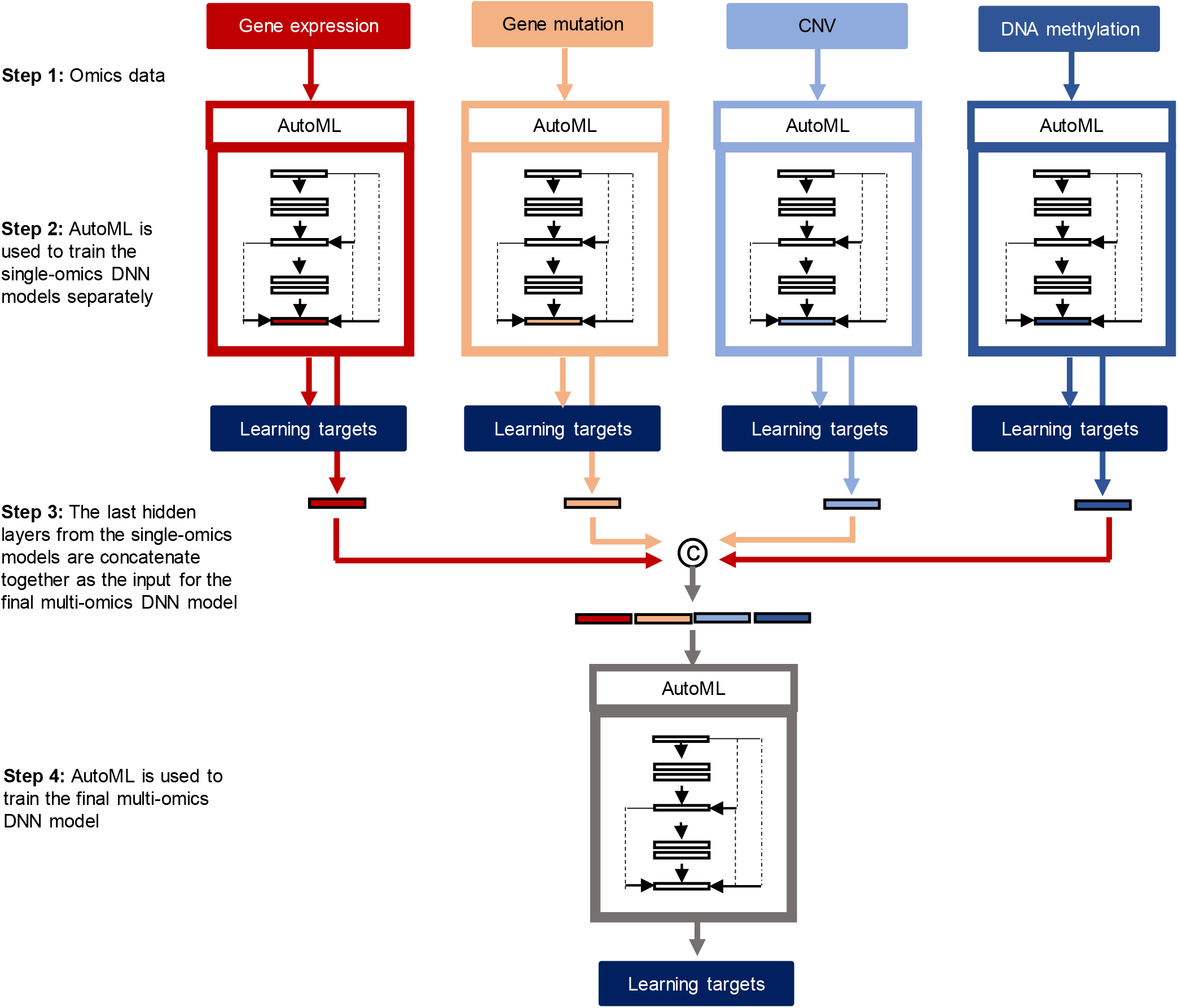
Scheme of AutoGenome for automatic multi-omics integration for AI model construction. **Step 1**: Collecting multi-omics data. Take multi-omics data of 4 data types as example, a gene expression matrix, a gene mutation matrix, a copy number variation (CNV) matrix and a DNA methylation matrix. **Step 2**: Use each single-type omics data respectively as input to train a model for the learning targets. Four network structure classes (MLP, ResNet, DenseNet and ENAS), numbers of layers, numbers of neurons per layer and hyperparameter combinations (batch size, learning rate and optimizers) was performed to search for a model with the highest performance evaluation scores as optimal single-omics models for this single-type omics data. Then, weights of the all optimal models were fixed, and stopped updating in the following steps. **Step 3**: Concatenate latent layers from all the optimal single-omics models. Here we chose all the last latent layers and extract the corresponding vector values into a concatenated vector as the input for **Step 4**. **Step 4**: Use the concatenated vector as input to train a model for the learning targets again. Same to the process in **Step 2**, search for optimal multi-omics model. Then, weights of this optimal model was fixed. All the optimal single-omics (from **Step 2**) and multi-omics (from **Step 4**) models were combined together into a whole network, as the final AutoGenome-based AI model.

To help researchers investigated the DNN model and learn about the impacts of each gene toward each learning target, we implement SHapley Additive exPlanations (SHAP)^13^ – a popular model explanation module into AutoGenome, which calculates marginal contribution of each feature to overall predictions, and summarized as a SHAP value, indicating potential feature importance to final biological meanings of interest.

To systematically evaluate the performances of AutoGenome, we apply AutoGenome on four different multi-omics tasks: a. drug response prediction, b. gene dependency prediction, c. breast cancer subtype prediction and d. pan-cancer patient stratification, the performances are compared both with popular multi-omics approaches and single-omics approaches, the results show us the significantly improvements of AutoGenome against the other approaches, and indicating AutoGenome to be a promising tool in multi-omics studies.

### Drug response prediction

New complex diseases arise along with changes in lifestyles and environment, creating new challenges and demands for new biomedicine treatments^14–16^. Although billions of dollars and tens of years have been spent on per de novo drug R&D, the success rate remains quite low. The main reason comes from safety issues and unclear mechanisms of actions for new drug candidates^14^. Drug repositioning, discovering new uses for existing FDA-approved drugs, can avoid the safety issues and skip toxicity testing, shorten time cost in R&D and increase success rate^15^. Famous examples e.g. Sildenafil initially for pulmonary arterial hypertension treatment is later found to treat erectile dysfunction^17^.

Large scale screen projects for cell lines under drug treatments offer rich resources for drug repositioning. Genomics of Drug Sensitivity in Cancers (GDSC)^18^ database measures half maximal inhibitory concentration (IC50). IC50 represented drug sensitivity scores for ~900 cancer cell lines under each of 265 anticancer drug treatments. GDSC also provides basal gene expression, gene mutation, DNA methylation etc., multi-omics data for the ~900 cell lines before drug treatments, including basal gene expression, gene mutation, DNA methylation etc. Based on GDSC, we aim to build an anticancer drug sensitivity prediction model. We take the gene expression and mutation profiles as features, and the IC50 values of 265 drugs as learning targets for each cell line, and run AutoGenome.

We implemented the optimal modelling by scanning structures of MLP, RFCN-ResNet and RFCN-DenseNet with various layer and neuron number setting, and found that RFCN-DenseNet-based networks achieved best performance for both gene expression and mutation single-omics models than other structures. Both of these two networks harbor single dense blocks (Figure 3B) with a net growth rate of 128 and 512 respectively; for the optimal multi-omics model part, a direct FC (Figure 3B) of MLP rather than RFCN-ResNet and RFCN-DenseNet was searched to be the best option (Figure 3A). Search strategy and search space are summarized in Methods Section. The above three searched models in total comprised the anticancer drug sensitivity prediction model.

**Figure 3.**
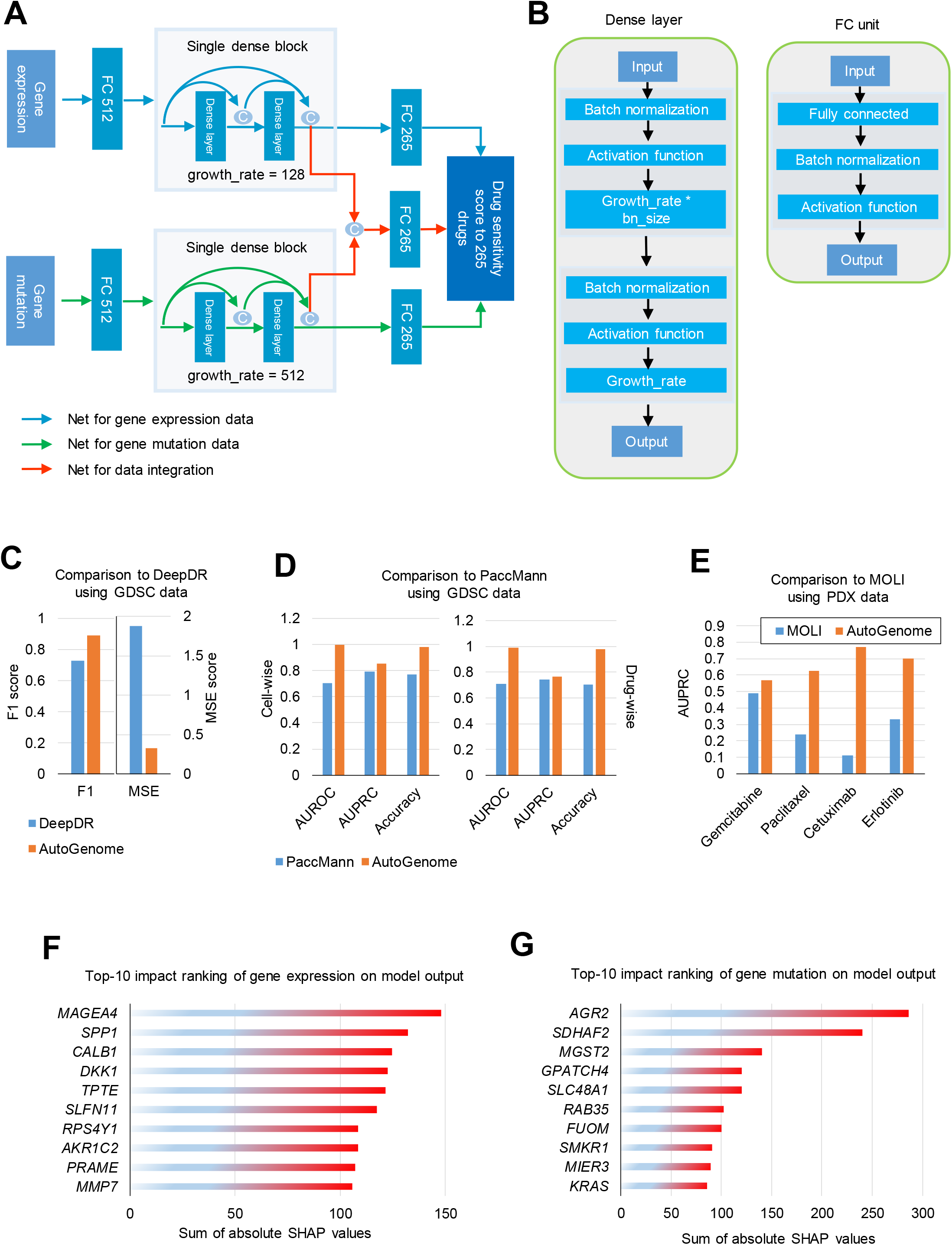
AutoGenome-based drug response prediction model. (A) Illustration of AutoGenome-based drug sensitivity prediction model using GDSC cancer cell line gene expression and mutation profiles to predict sensitivity scores to 265 anticancer drugs. (B) Illustration of Automics basic net units – dense layer units for DenseNet and FC units. (C) Comparison of AutoGenome based drug sensitivity model to DeepDR using GDSC data. GDSC IC50 values were used as golden-standard positives. F1 score and MSE score were used for evaluation comparison. (D) Comparison of AutoGenome based drug sensitivity model to PaccMann using GDSC data. We used SMILES of 28 drugs and 936 cell lines shared in training sets of both our model and PaccMann for comparison. GDSC IC50 values were used as golden-standard positives. Evaluation comparison were performed in both cell-wise and drug-wise. (E) Independent validation of AutoGenome based drug sensitivity model in a PDX mice data set and comparison to MOLI. Independent validation of AutoGenome in a mice PDX data set. We tested the GDSC-IC50-trained AutoGenome model on a mice PDX data, which uses tumor size reduction as learning targets. Here tumor size reduction quantity was used as ground truths for performance evaluation. 4 drugs shared between our model and MOLI were used for comparison. (F) Top-10 gene expression ranked by impact on model output for AutoGenome based drug sensitivity model. (G) Top-10 gene mutation ranked by impact on model output for AutoGenome based drug sensitivity model.

Mean squared error (MSE) and spearman correlation coefficient (SCC) of the optimal single-omics models of gene expression were 1.532 and 0.8717, and 1.9 and 0.8463 for gene mutation model. MSE and SCC of the optimal multi-omics model were obviously improved, with an obviously decreasing MSE 0.3266 and increasing SCC 0.967 (Figure 3C). We utilized log-transformed IC50 < −2 (approximately 0.135 μM) as standard threshold^19^ to define drug sensitive (positive) and resistant (negative) groups, and found that the multi-omics model also outperformed single-omics models in area under the receiver operating characteristic curve (AUROC), precision, recall and accuracy (Supplementary Figure 3A). Our results demonstrate that there is a significant improvement by multi-omics data integration using AutoGenome than only using single omics data for deep learning modelling.

We compared the AutoGenome based drug response prediction model with popular existing models - DeepDR^6^, PaccMann^5^ and MOLI^8^. DeepDR reports MSE as 1.96 in the original paper^6^. We reproduced the network architecture of DeepDR and achieved a MSE as 1.8793 and F1 as 0.7283 using the same data for the AutoGenome, which is significantly outperformed by the AutoGenome (MSE 0.3266, F1 0.8907, Figure 3C). For PaccMann, we randomly queried 28 drugs for IC50 predictions from its webserver to compare with AutoGenome predictions, and found that AutoGenome showed higher AUROC, AUPRC (area under precision recall curve) and accuracy in both cell-wise (AUROC 0.998 vs. 0.702, AUPROC 0.854 vs. 0.792, accuracy 0.982 vs. 0.768) and drug-wise (AUROC 0.987 vs. 0.708, AUPROC 0.764 vs. 0.742, accuracy 0.978 vs. 0.705) levels (Figure 3D). For MOLI, it trained response prediction models for 4 drugs (Paclitaxel, Gemcitabine, Cetuximab, Erlotinib) using GDSC data, and evaluated in a patient-derived xenograft (PDX) mice data set^20^. The PDX data includes gene expression, mutation and copy number variation (CNV) profiles for 399 mice and tumor size reduction as drug response index to 63 drug treatments for each mouse. To compare with MOLI, we predicted responses for the 4 drugs by inputting the PDX mice gene expression and mutation into the AutoGenome model and fetching corresponding predictions for the 4 drugs from total 265 drugs. The results showed that AutoGenome achieved higher AUPRC value for the 4 drugs than MOLI (Paclitaxel 0.616 vs. 0.24, Gemcitabine 0.558 vs. 0.49, Cetuximab 0.771 vs. 0.11, Erlotinib 0.7 vs. 0.33, Figure 3E).

In addition, we compared AutoGenome with other strategies of omics data integration: concatenated raw input + MLP (F1: 0.707), VAE latent + MLP (F1: 0.728), VAE recon + MLP ((F1: 0.733), and raw input + AutoGenome (F1: 0.725), gene expression + AutoGenome (F1: 0.7315) and gene mutation + AutoGenome (F1: 0.708), and we found that AutoGenome showed the highest F1 score as 0.891 among all using the same data sets (Supplementary Figure 3B). Interestingly, the F1 scores were sorted as: AutoGenome > AutoGenome for gene expression > AutoGenome for gene expression and mutation concatenation > AutoGenome for gene mutation (F1: 0.8907 > 0.7315 > 0.7253 > 0.7075). It indicates that integrating multi-omics raw input data for modelling directly caused neutralization of good-performance data (gene expression, F1: 0.7315) and poorly-performance data (gene mutation, F1: 0.7075), leading to worse predictions (F1: 0.7253) than that of the good-performance one; But in contrast, AutoGenome largely promoted the performance for multi-omics data integration (F1: 0.8907) better than each single data (Figure 3D).

When analyzing importance of features contributing to the final prediction, we listed top gene expressions and gene mutations ranked by SHAP values that showed highest importance to all 265 drug response predictions (Figure 3E and 4F). Functional enrichment analysis showed that top-50 ranked gene expressions were enriched in doxorubicin or daunorubicin metabolic process, regulations of cell proliferation, growth factor activity and Wnt signaling pathway etc.; and the top-50 ranked gene mutations were enriched in regulations of gene expression, cell proliferation and apoptotic process and typical signal pathway in cancers. Genes of the expression features enriched in location of extracellular space and exosome. Interestingly, unlike gene expression, the mutation feature genes were enriched in location of mitochondrion and endoplasmic reticulum.

**Figure 4.**
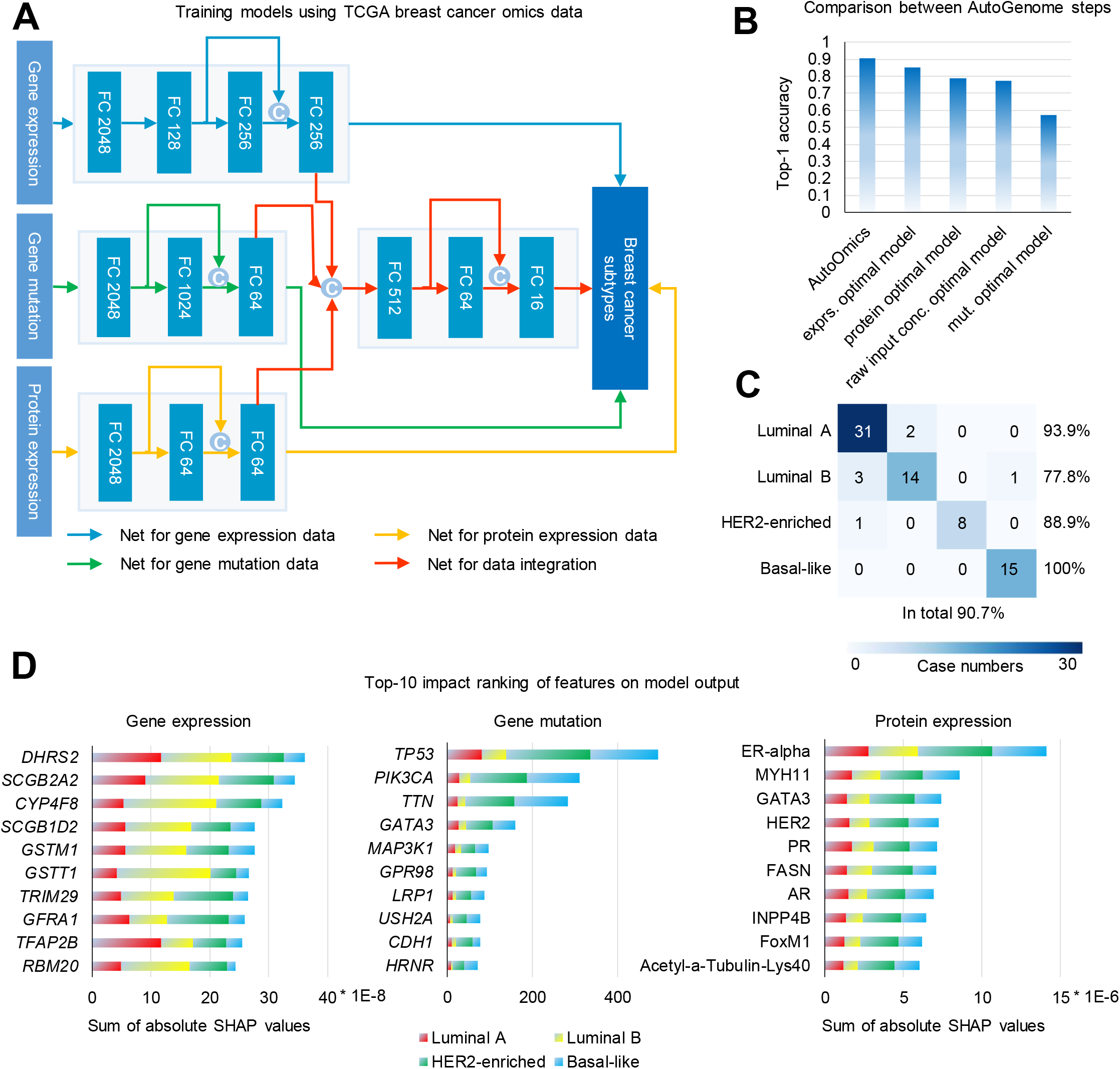
AutoGenome-based cancer subtyping and pancancer type prediction models. (A) Illustration of AutoGenome-based cancer subtyping prediction model using TCGA breast cancer patient gene expression, gene mutation and protein expression profiles to predict four breast cancer subtypes. (B) Performance comparison between AutoGenome steps using top-1 accuracy. Total data were randomly separated in 7:2:1 ratio for training, evaluation and test datasets. The test dataset was used for top-1 accuracy calculation. (C) Confusion matrix using predictions of AutoGenome-based cancer subtyping prediction model. True labels are in the row and predictions are in the column. Top-1 accuracy for each of the four breast cancer subtypes and total top-1 accuracy are indicated. (D) Feature importance analysis. Top-10 features ranked by impact on model output are performed for gene expression (left), gene mutation (middle) and protein expression (right).

Besides, we tested AutoGenome on gene dependency prediction. With the help of single-gene perturbation techniques e.g. RNA inference (RNAi) and CRISPR gene editing, researchers can perturb genetics to study effect of single gene to diseases or biological processes (e.g. by measuring mortality rate of cancer cells following by a gene RNAi)^21^. Through integrating basal multi-omics data of samples of interest before gene perturbations and linking to learning targets of phenotype data after gene perturbation, researchers can train a gene dependency model to predict which genes play essential roles for the phenotype. In this way, the researchers don’t need to actual perturbation on that genes, and it aids in large-scale screening for identifying gene targets. The Cancer Dependency Map (DepMap)^21^ includes CRISPR screen results on 17635 genes for cancer cell lines and corresponding basal omics datasets for each cell. To focus on cancer causal genes, we narrowed down the screen results to 610 studied pan-cancer or cancer-type-specific priority gene targets^22^, and used gene expression and mutation profiles as features to run AutoGenome. AutoGenome firstly scanned network structures of MLP, RFCN-ResNet and RFCN-DenseNet with different numbers of layers and neurons, and outputted RFCN-DenseNet-based structures with 4 and 5 dense blocks, with net growth rates as 16 and 8 respectively, as the best models for gene expression and mutation profiles (Supplementary Figure 4A), which achieved MSE as 0.04011 and 0.04495, and SCC as 0.8610 and 0.8521 respectively. Then AutoGenome scanned best net structures following concatenating the last hidden layers of the above two models, and returned a three-layer MLP with neuron numbers of 32, 32 and 610 (Supplementary Figure 4A), which achieved a better MSE and SCC as 0.03428 and 0.8777 (Supplementary Figure 4B). Gene dependency score < −0.6 standard threshold^21^ were set to define gene perturbation sensitive (positive) and resistant (negative) groups. And the multi-omics model also outperformed single-omics models in five classification evaluation indexes, which demonstrates improvement by multi-omics data integration using AutoGenome for the task than only using single omics data. For performance comparison, we constructed trials same to that of drug sensitivity prediction, and it showed AutoGenome-based model outperformed all other methods for gene dependency prediction task (Supplementary Figure Figure 4C).

We also listed top gene expressions and gene mutations ranked by SHAP values that showed highest importance to all 610 gene dependency predictions (Supplementary Figure 4D and 4E). Functional enrichment analysis showed that top-50 ranked gene expressions were enriched in regulations in signal transduction and cell proliferation, immune response and renin-angiotensin system etc, and the location was enriched in extracellular exosome and space. The top-50 mutation feature genes were enriched in location of cortical cytoskeleton.

### Breast cancer subtype prediction

Large-scale accumulation of multi-omics data and electronic medical records for patient individuals makes it possible to study precision medicine, that is, to specify medicine treatments for each patient according to their personal clinical responses and physiological and genomics features. These features may include susceptibility to diseases, mechanisms of onsets, prognosis conditions, responses to specific treatments and genetics background etc^23^. For complex diseases e.g. cancer, although patients may belong to the same cancer types based on pathology, their response to drugs or immune therapy often vary largely. This is may be due to theirs difference in genetics background^24^. Therefore, it is necessary to take advantage of patients’ multi-omics data for cancer subtyping.

The Cancer Genome Atlas (TCGA) database includes six types of omics data in patient individual level for more than 20 cancer types^25^. Here we implemented AutoGenome to build a classification model for breast cancer subtype prediction using patients’ gene expression, gene mutation and protein expression profiles as features, and focused on four subtypes of PAM50-profiling-test based, luminal A, luminal B, triple-negative and HER2-enriched^26^. Apart from searching optimal models using MLP, RFCN-ResNet and RFCN-DenseNet structures, we also included ENAS randomly-generated structures for this classification task, and found it showed outstanding performance than RFCN-DenseNet/ResNet. For optimal single-omics model of gene expression, AutoGenome returned an ENAS-based model containing 4 FC layers with 2048, 128, 256 and 256 neurons per layer and one skip connection concatenating the 128- and 256-neuron layer outputs as input of the 256-neuron layer (Figure 5A). It achieve top-1 accuracy 0.8533 for breast cancer subtyping (Figure 5B), better than published graph deep learning based model using the same origin of dataset and learning targets (with an accuracy of 0.8319)^27^. For optimal single-omics models of gene mutation and protein expression, AutoGenome returned both ENAS models with three layers and one skip connections (Figure 5A), showing top-1 accuracy as 0.573 and 0.787 respectively, lower than that of gene expression (Figure 5B). Then AutoGenome linked all the three models and again generated an optimal model for data integration as an ENAS-based three-FC-layer network with 512, 64 and 16 neurons (Figure 5A), with an improved top-1 accuracy as 0.907 (Luminal A: 0.939, Luminal B: 0.778, HER2-enriched: 0.889, Basal-like: 1), significantly better than each of the single-omics model (Figure 5B and 5C). It outperformed a published approach using SMO-MKL, which achieved a 0.798 average accuracy of any two immunohistochemistry-marker-based subtypes^28^. When direct concatenating three omics data together, the top-1 accuracy is 0.773, better than that of gene mutation and worse than gene expression and protein expression (Figure 5B).

Then we analyzed importance scores represented by SHAP values for each feature to the model outputs. SHAP value in gene expression showed that *RBM20* gene expression contributed more in luminal B subtype than other subtypes (Figure 5D). The finding is accordant with the public results which proves that *RBM20* gene expression is correlated with PDCD4-AS1/PDCD4, a tumor suppressor in TNBC cell lines^29^ and a subset of TNBC potentially benefit therapy targeting luminal subtype’s typical pathways^30^. In other examples, the SHAP value of *TFAP2B* shows it is an important gene feature in breast cancer (Figure 5D). It is proven by the HMAN PROTEIN ATLAS which shows *TFAP2B* (ENSG00000008196) is a cancer-related gene and its expression is highest in breast cancer pathology data and enriched in breast cancer. Besides, SHAP value of *TFAP2B* is highest in luminal A, which is agreement with that *TFAP2B* is associated with WNT/β-catenin pathway in luminal breast cancer, and its encoding protein AP-2 transcription factor regulates luminal breast cancer genes^31^.

In protein expression level, based on SHAP values, ER-alpha ranked top 1 among all protein expression and showed important contribution to all four subtype in breast cancer (Figure 5D). This phenomenon also is accordance with the research of estrogen receptors which is a very important marker for prognosis and a marker that is predictive of response to endocrine therapy in breast cancer^32^. The loss of ER expression portends a poor prognosis and, in a significant fraction of breast cancers, this repression is a result of the hypermethylation of CpG islands within the ER-alpha^33^.

To systematically analyze the reliability of feature importance results, we used SHAP-ranking top features to represent samples and checked inner- and intra-subtype similarity of samples. It showed that compared to using all features representing samples, SHAP-top-ranked features could better cluster samples of the same subtypes together and distinguish between intra subtypes, which was quantified by silhouette score (Supplementary Figure 4). The scores achieved highest using top features (top 70, 40 and 10 for gene expression, gene mutation and protein expression), and then decreased when more bottom-ranked features were included (Supplementary Figure 4). It demonstrates that top-ranked features by SHAP values show the direct role to improve breast cancer subtype classification.

Additionally, we built a 24 cancer type prediction model using TCGA pancancer omics data. AutoGenome trained a 6-FC-layer and 4-skip-connection ENAS network for gene expression profiles with top-1 accuracy 0.963, and a 4-FC-layer and 2-skip-connection ENAS network for gene mutation profiles with top-1 accuracy 0.681 (Supplementary Figure 9A and 9B). AutoGenome linked the two networks using a 3-FC-layer and 1-skip-connection ENAS network, achieving an improved top-1 accuracy 0.973 (Supplementary Figure 9A and 9B). For comparison, we also performed stacked ensemble learning to link the two networks by using their softmax target layers rather than last hidden layers as a new network input. We tested both ENAS and MLP for the ensemble learning way, and the top-1 accuracy was consistently round 0.857, which lied in between that of gene expression ENAS network and the gene mutation ENAS one, lower than that of AutoGenome (Supplementary Figure 9B). It demonstrated that AutoGenome outperformed stacked ensemble learning in this task.

## Discussion

Genomics data are widely accumulated using high-throughput sequencing for cell lines, animal models and patient individuals. Different omics data types can reflect different aspects of features for samples, thus it is important to integrate all omics data for one sample at the same time. To address it, we developed a new version with multimodal data integration function for – AutoGenome. AutoGenome firstly builds single-omics models for each single data, and the latent spaces extracted from single-omics models were combined together and then are subjected to deep learning modeling for the same learning target as single-omics.

Weights of single-omics models are frozen once finished training. They are no longer updated in the stage of multi-omics modelling using combined latents. Unlike AutoGenome, published methods treat each of single-omics data as a branch of networks (subnetworks)^5^, that is to say, weights of certain subnetwork are dependent on other’s subnetworks, and they are kept updating simultaneously following the whole network training. This difference may be the reason why AutoGenome is outstanding for efficient data integration. AutoGenome may maximally extract latent that can best represent learning targets for each data individually. Our three prediction tasks also demonstrate the excellent performance of AutoGenome over other methods.

Our data integration idea is similar to stacking ensemble learning^34^ but not the same. Stacking ensemble emphasizes combining predictions of individual models not the latent spaces as we have proposed.. Furthermore, traditional methods take all the multi-omics data as a whole input for a model, but in this paper, we demonstrate that taking each of single-omics data to build independent models and then performing combination as a whole are better choices. The effectiveness of our method is probably because that batch effect, difference in source of data and value ranges among data cannot be well removed by normalization. Direct combining raw values of different data types will lead to unexpected bias, thus degrade performance of the model (Figure 3D, Figure 4C and Figure 5B).

As we have claimed on AutoGenome^10^, CNN and RNN are not suitable to modeling genomics profiles because features of genomics data are non-Euclidean, thus pure FC is better. Published methods mainly utilize MLP, where layers only link neighboring layers^4–6^. It is not efficient to model biological regulations and feedback loops between different levels. Thus we introduced skip connections in our network design. We tried RFCN-ResNet and RFCN-DenseNet – two classical network structures with skip connections. In our experiments, RFCN-DenseNet was proved to be better than RFCN-ResNet in both drug sensitivity and gene dependency tasks. This is may be due to that densly skip-connections in RFCN-DenseNet may cover more possible interaction combinations than RFCN-ResNet. Interestingly, RFCN-DenseNet for drug sensitivity is wider (with more number of neurons per layer) and shallower (with less number of layers) than that for gene dependency task, no matter using gene expression or mutation data (Figure 3A and Figure 4A), which may imply that these anticancer drugs can influence more targets and pathways initially than single gene perturbations, but the latter one can expand deeply in biological network. However RFCN-DenseNet cannot keep good performance in cancer subtyping classification task, both RFCN-ResNet and RFCN-DenseNet showed low accuracy and ENAS-based randomly-generated network significantly outperformed them. It imply that ENAS can achieve a larger searchable space than RFCN-DenseNet, where the structure of the latter one can only be extended in a fixed manner.

Taking all above, AutoGenome is the first automated machine learning tool for multi-omics research. It comprises novel network units and network architectures specificly designed for genomics data. And it also integrate built-in novel multimodal integration method for efficiently taking advantages of multi-omics data. Besides, AutoGenome also will provide more biological explanation for predicted results by sharp, which can help biologist to discovery more interesting biological marker to do research. AutoGenome surely speed up bioinformatics and genomics study and aid in dissecting important findings for biological researchers.

## Methods

### Hyper-parameter search

Hyper-parameter search method refers to our previous described approach in AutoGenome^10^. The hyper-parameters in search space are learning rate, total batch size, momentum, weight decay, number of layers in neural networks and number of neurons in each layers.

#### RFCN-ResNet Search Space

Search space are as followings. 1) The number of blocks for ResNet, default value is [1, 2, 3, 5, 6]. 2) The number of neuros in each layer, default value is [8, 16, 32, 64, 128, 256, 512, 1024, 2048]. 3) The drop-out ratio of the first layer compared with the input layer, selected from [0.6, 0.8, 1.0]

#### RFCN-DenseNet Search Space

Search space are as followings. 1) The blocks structure for DenseNet, default value is [[2, 3, 4],[3, 4, 5]]. 2) The growth rate of neuros in each block, default value is [8, 16, 32, 64, 128, 256, 512, 1024, 2048]. 3) The drop-out ratio of the first layer compared with the input layer, selected from [0.6, 0.8, 1.0].

### Efficient Neural Architecture Search

ENAS search method refers to our previous described approach in AutoGenome^10^. Search space are as followings. 1) The number of neurons in from the 2rd layer to the last layer, selected from [16, 32, 64, 128, 256, 512, 1024, 2048]. 2) The connection relationship between different layers.

### Data collection and preprocessing

For drug sensitivity and gene dependency prediction tasks, we downloaded gene expression and mutation data, drug response data and CRISPR-based gene dependency data from GDSC (https://www.cancerrxgene.org) and DepMap (https://depmap.org/portal/download/) web resources. Gene expression profile includes 1018 cancer cell lines and 17418 genes. Values were log2-transformed. Gene mutation data covers 974 cancer cell lines. We set discrete values of 1 and 0 indicating somatic mutated or wide-type status and removed silent mutation cases, remaining union set of 19350 genes for analysis. Drug response data includes log-transformed IC50 values for 990 cancer cell lines representing response to 265 single anticancer drug treatments. K-nearest-neighbor algorithms was performed to fill in missing values for the IC50 response data by R function knn. Gene dependency data covers 558 cancer cell lines and response to CRISPR perturbation of 17634 human genes. To focus on cancer priority gene targets^22^, 610 gene CRISPR cases were remained for analysis. Cell lines were mapped using identifiers of Catalogue Of Somatic Mutations In Cancer (COSMIC) between datasets, thus remained 936 cell lines for drug sensitivity task and 324 for gene dependency task.

For cancer subtyping prediction task, breast cancer patients’ gene expression, mutation and protein expression data were downloaded from TCGA (https://gdac.broadinstitute.org/), where feature numbers were 20531 genes, 16806 somatic mutated genes and 226 proteins respectively. Gene expression values were log2-transformed and silent mutations were removed from gene mutation data. PAM50-based subtypes for patients were downloaded from published paper^35^. 396 patients shared between the feature data and subtype data were used for cancer subtyping prediction task. For pancancer type prediction, 5,780 patient samples with gene expression and somatic mutated gene profiles were used for modelling.

### Model training, evaluation and explanation

For drug sensitivity and gene dependency tasks, data were randomly separated into 8:1:1 ratio for training, evaluation and test data. Evaluation was performed based on 10-fold cross validation to calculate MSE and SCC. Precision, recall, accuracy and AUROC were calculated when using a threshold to group positives and negatives. For cancer subtyping task, the ratio was set as 6:2:2 in ENAS-based modelling, since evaluation data was used to update and determine net architectures for ENAS. Data were splitted in a stratified manner to make percentage of subtypes equal between data sets. Top-1 accuracy for the test data was used as final evaluation. All evaluation scores were calculated using python sklearn module. Model explanation is performed by “SHAP” module implemented within AutoGenome. AutoGenome take the best model and raw data as input, when calling with “autogenome.explain()”. And then AutoGenome will automatically return the SHAP value of each feature for each sample for further interpretation.

### Software Availability

We will open the utilization of AutoGenome package to the public upon the acceptance of manuscript.

## Acknowledgements

We thank Prof. Mingyue Zheng for critical reading and suggestions for revision.

## Author Contributions

N.Q. designed and conceived the project. C.X., D.L., L.Z., Z.X., W.H. and D.W. implemented the AutoGenome V2 python package and perform data experiment under the guidance of N.Q. C.X., D.L., L.Z and N.Q. wrote the paper. N.Q. and M.Z. revised the manuscript. All authors read and approved the final manuscript.

## Competing Interests statement

The authors declare no competing interests.

---

**Supplementary Figure 1.**
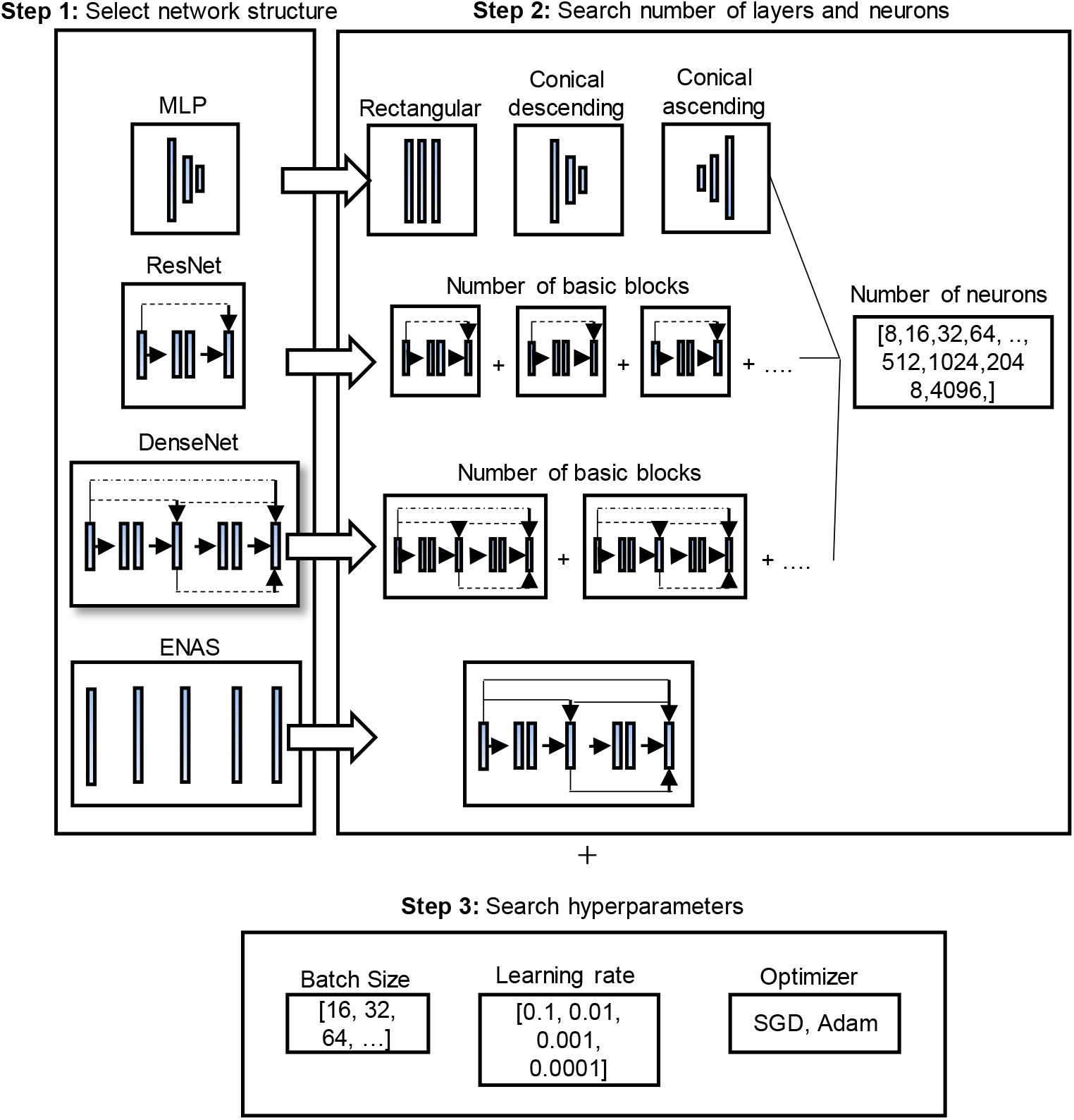
Process of searching for optimal model structures and hyperparameters (Related to Figure 2).

**Supplementary Figure 2.**
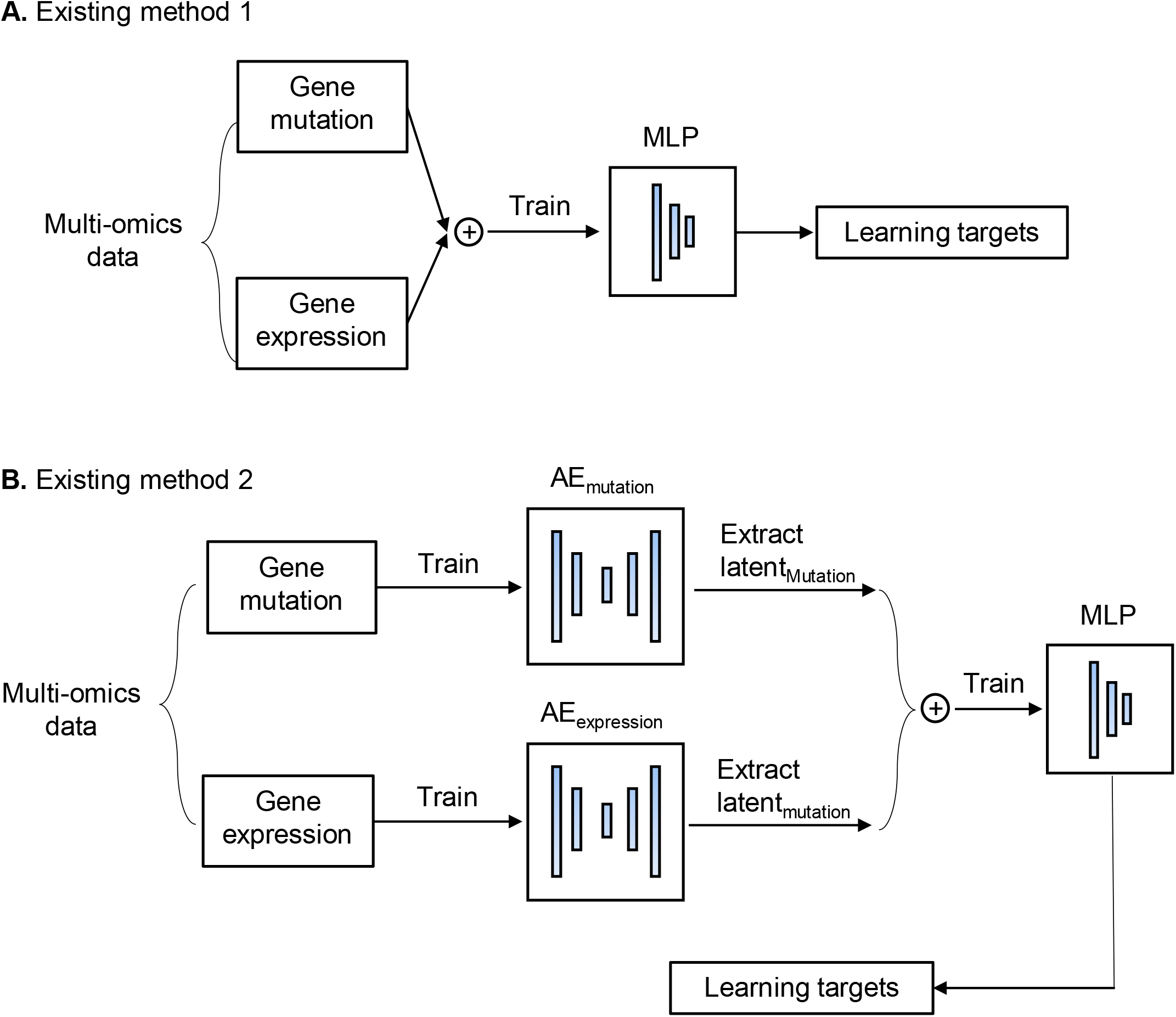
Existing methods for integrative analysis of multi-omics data (Related to Figure 3). Here we take multi-omics data of two data types (gene mutation matrix and gene expression matrix). **(A)** Existing method 1: Direct concatenate two data matrix together as input to train a MLP against learning targets. **(B)** Existing method 2: Firstly train two autoencoders (AEs) for gene mutation and gene expression respectively, then extract the output vectors of their encoders (latent_Mutation_ and latent_Mutation_) and concatenate together as input to train a MLP against the learning targets.

**Supplementary Figure 3.**
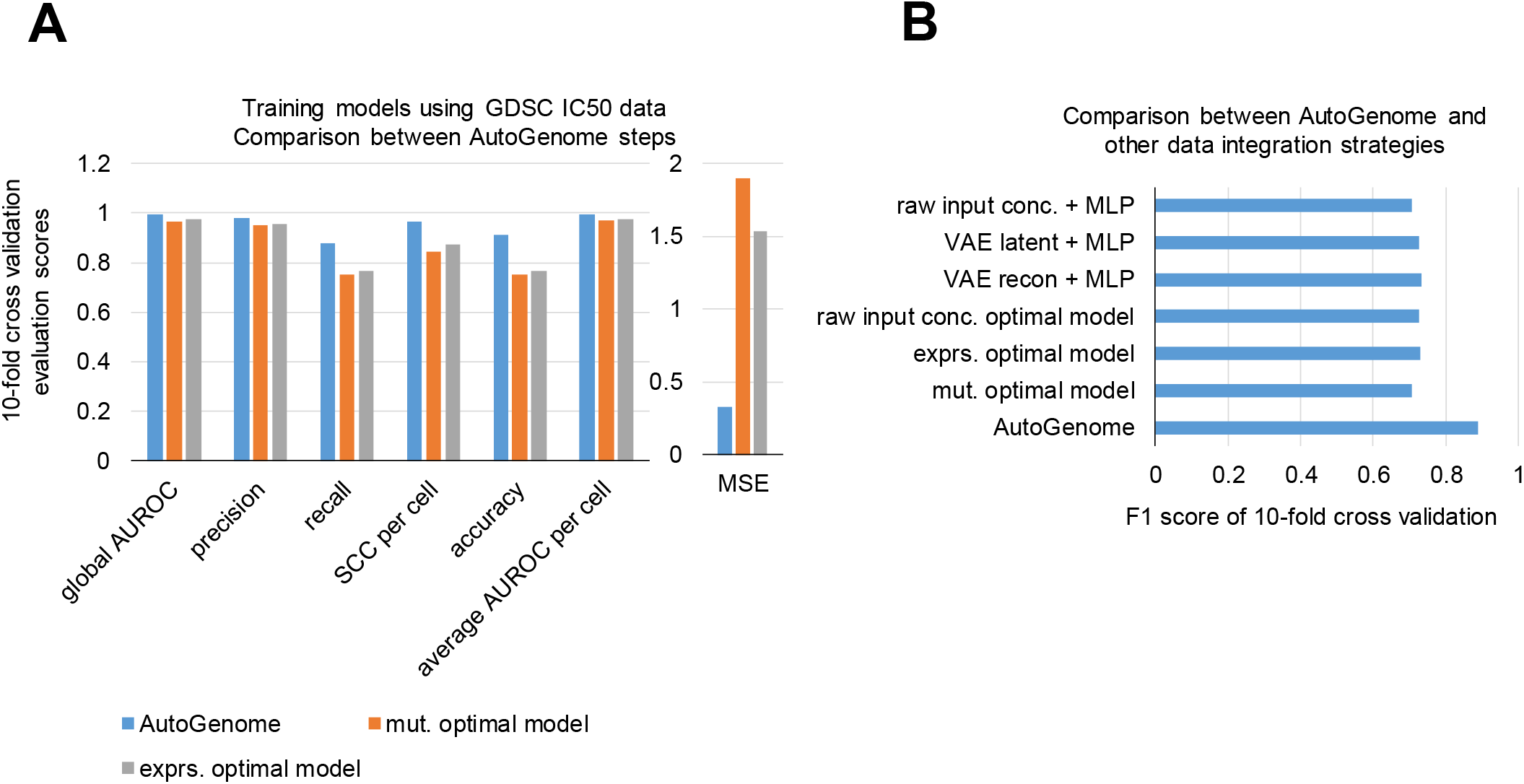
Comparison of AutoGenome-based drug response model to other methods (Related to Figure 3). **(A)** Performance comparison between AutoGenome steps using 7 evaluation scores. All scores were calculated in 10-fold cross validation manner using GDSC IC50 values as ground truths. For SCC and average AUROC per cell, values were firstly calculated for each cell line and then averaged across cell lines. For global AUROC, prediction matrix of cells vs. 265 drugs were expanded to a vector, then performed for calculation globally. **(B)** Performance comparison between AutoGenome and other data integration strategies using F1 score.

**Supplementary Figure 4.**
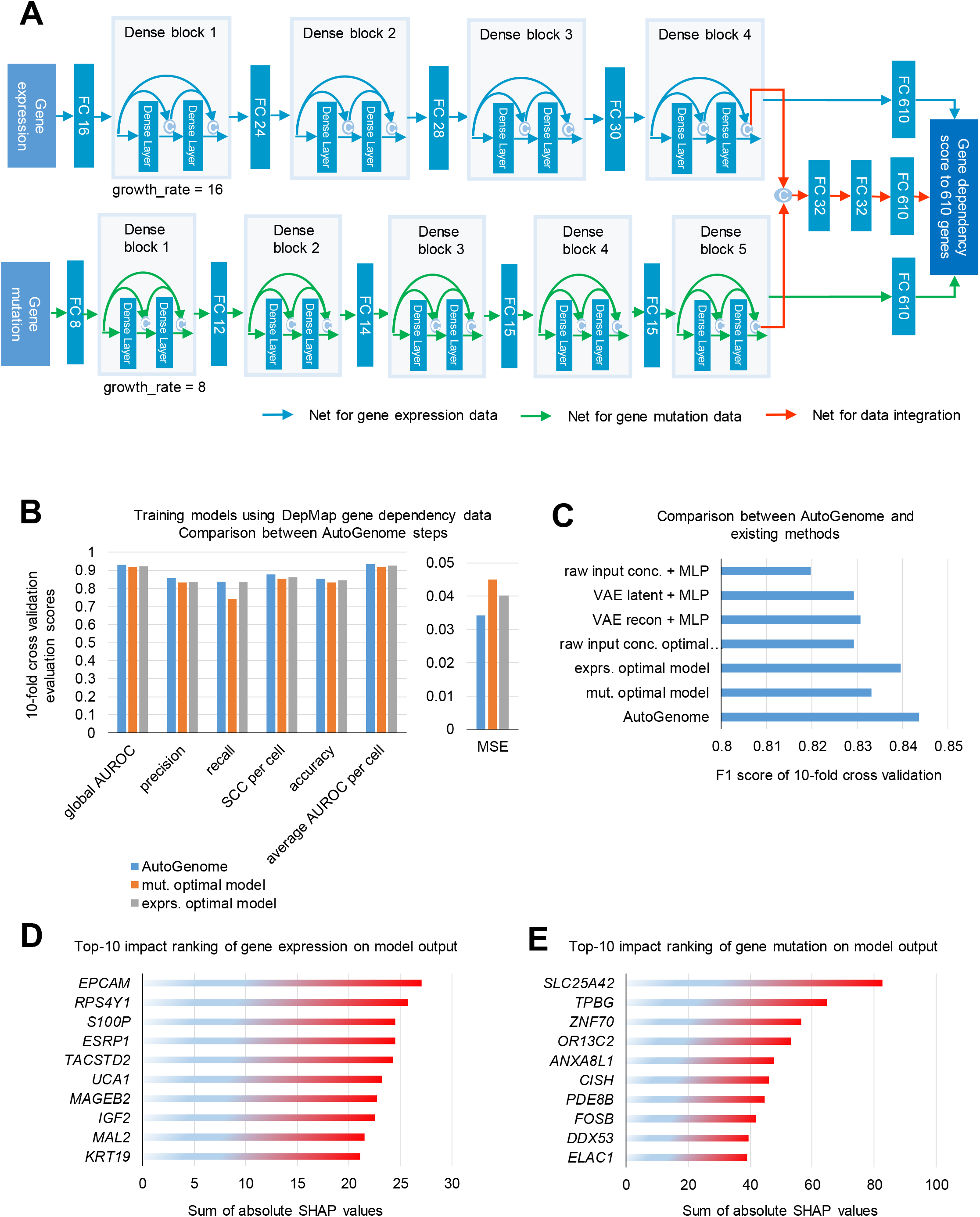
AutoGenome-based gene dependency prediction model (Related to Figure 3). (A) Illustration of AutoGenome-based gene dependecy prediction model using DepMap cancer cell line gene expression and mutation profiles to predict dependency scores to 610 cancer target genes. (B) Performance comparison between AutoGenome steps using 7 evaluation scores. All scores were calculated in 10-fold cross validation manner using DepMap gene dependency scores as ground truths. For SCC and average AUROC per cell, values were firstly calculated for each cell line and then averaged across cell lines. For global AUROC, prediction matrix of cells vs. 610 genes were expanded to a vector, then performed for calculation globally. (C) Performance comparison between AutoGenome and existing methods using F1 score. Feature importance analysis for AutoGenome are shown in (D) and (E). (D) Top-10 gene expression ranked by impact on model output. (E) Top-10 gene mutation ranked by impact on model output.

**Supplementary Figure 5.**
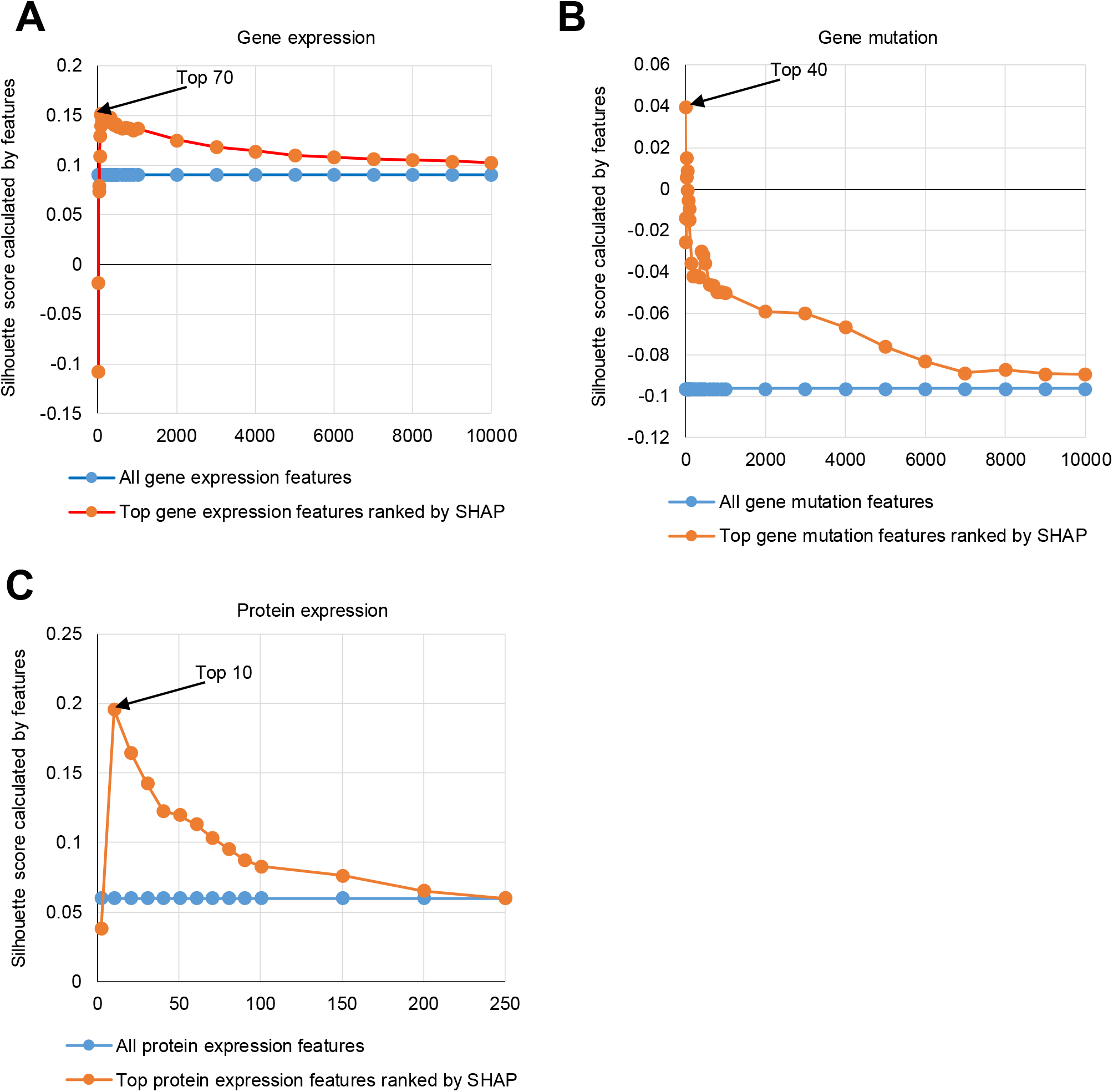
Silhouette scores calculated using top features ranked by SHAP values (Related to Figure 4). Silhouette scores of breast cancer patient samples to four subtyping classification calculated by top-ranked features by SHAP (orange) and all features (blue) of gene expression **(A)**, gene mutation **(B)** and protein expression **(C)**. Top 70, 40 and 10 features show the highest silhouette scores for gene expression, gene mutation and protein expression respectively.

**Supplementary Figure 6.**
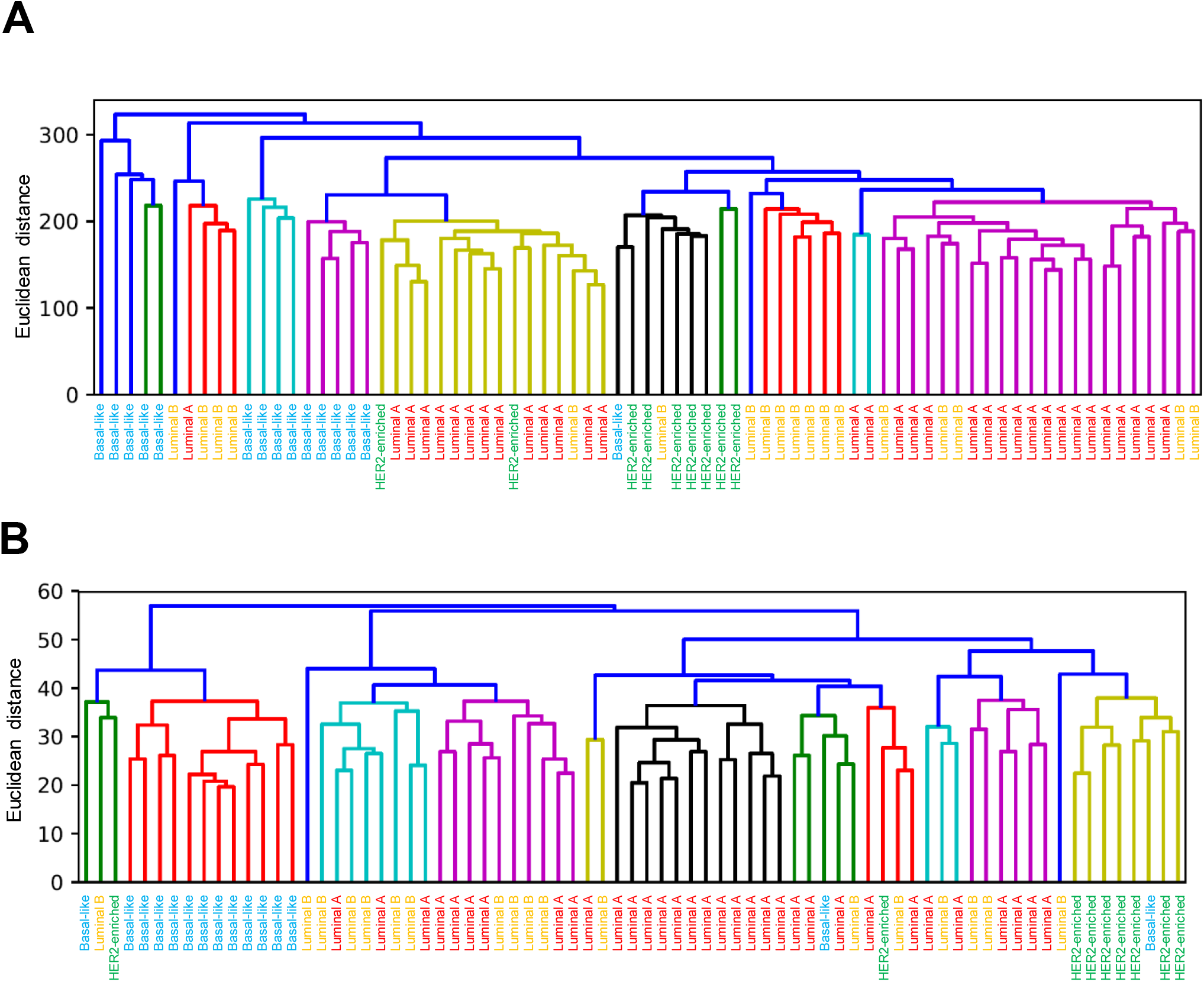
Hierarchical clustering of breast cancer samples using all gene expression features (A) and top-70-SHAP ranked gene expression features (B) (Related to Figure 4).

**Supplementary Figure 7.**
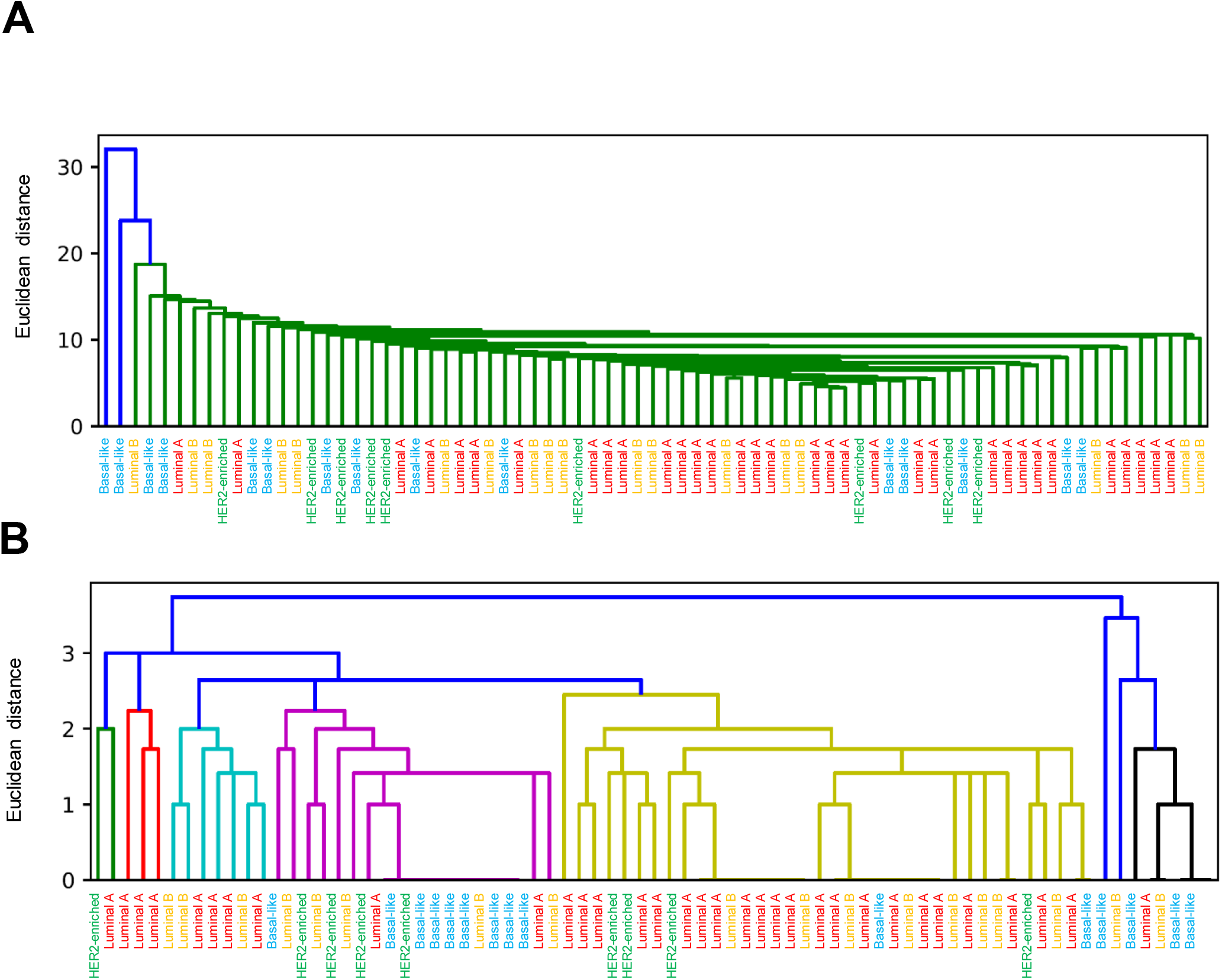
Hierarchical clustering of breast cancer samples using all gene mutation features (A) and top-40-SHAP ranked gene mutation features (B) (Related to Figure 4).

**Supplementary Figure 8.**
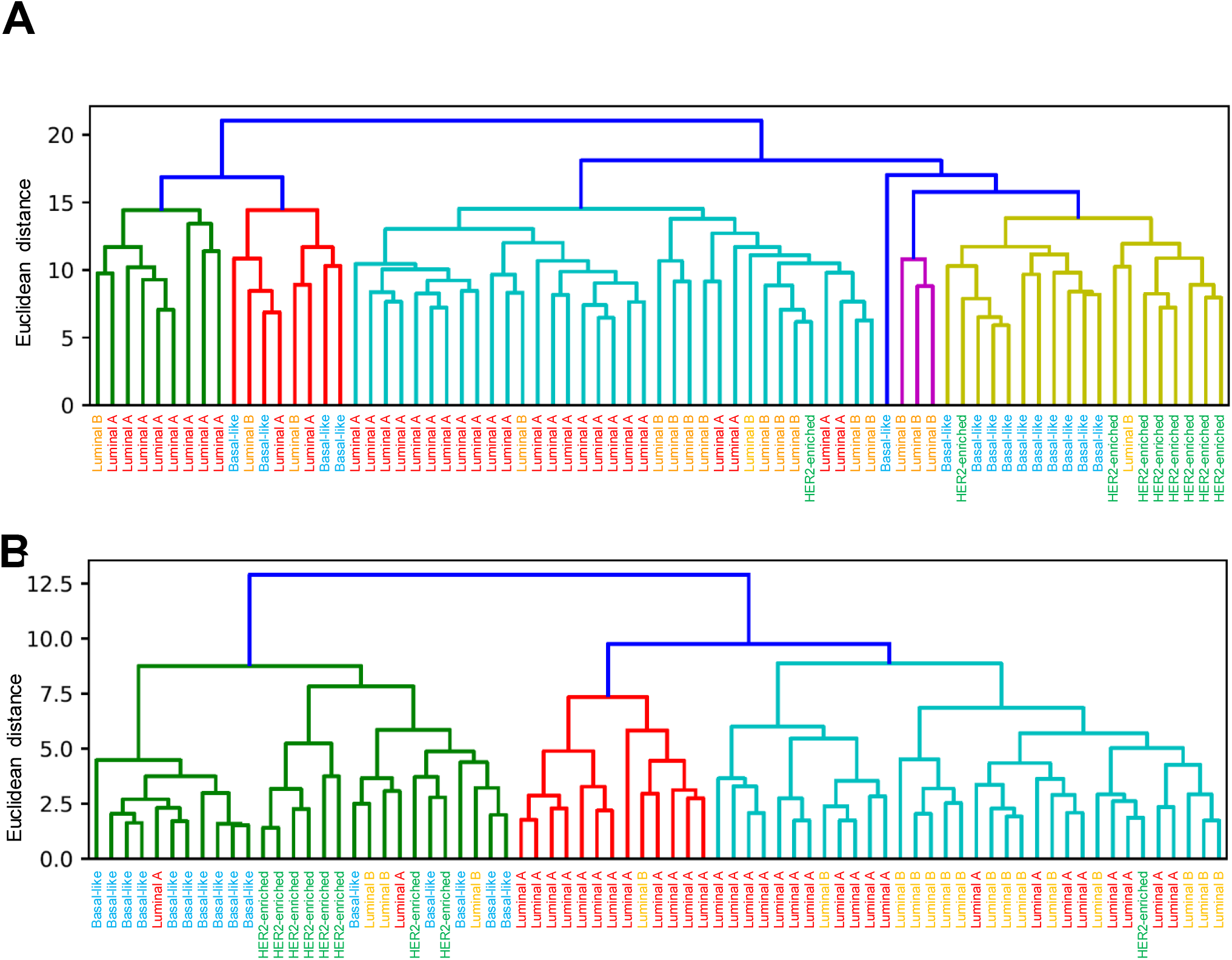
Hierarchical clustering of breast cancer samples using all protein expression features (A) and top-10-SHAP ranked protein expression features (B) (Related to Figure 4).

**Supplementary Figure 9.**
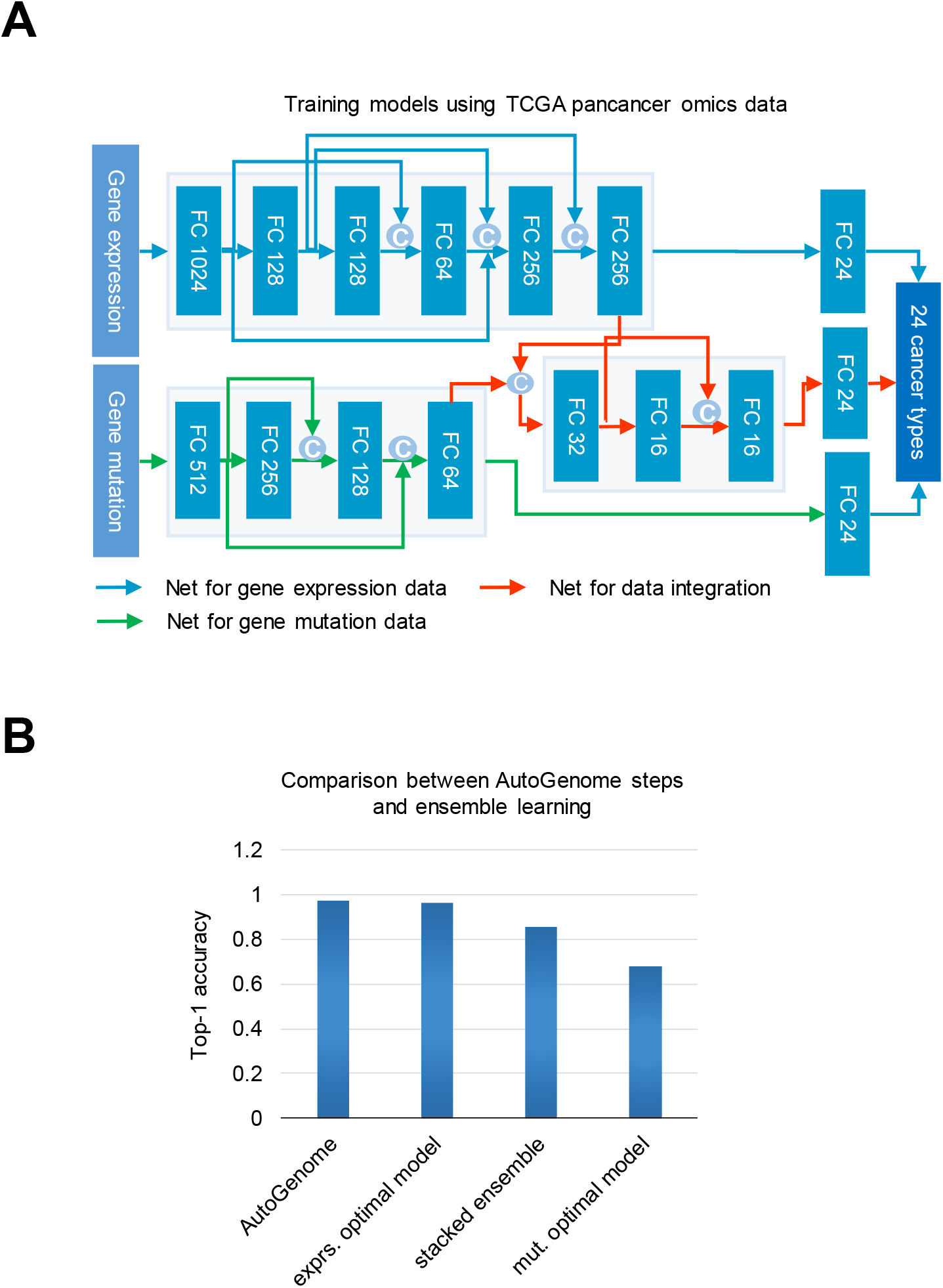
AutoGenome-based gene dependency prediction model (Related to Figure 4). (A) Illustration of AutoGenome-based cancer type prediction model using TCGA pancancer gene expression and gene mutation profiles to predict 24 cancer types. (B) Performance comparison between AutoGenome steps and ensemble learning using top-1 accuracy. Stacked ensemble learning was performed by concatenating softmax vectors of target prediction layers and to train ENAS or MLP prediction networks.

## Notes

### Competing Interest Statement

The authors have declared no competing interest.

## References

1. Yngvadottir, B., MacArthur, D. G., Jin, H. & Tyler-Smith, C. The promise and reality of personal genomics. Genome Biol. 10, 237 (2009).

2. Mayakonda, A. & Koeffler, H. P. Maftools: Efficient analysis, visualization and summarization of MAF files from large-scale cohort based cancer studies. bioRxiv 052662 (2016) doi:10.1101/052662.

3. Zou, J. et al. A primer on deep learning in genomics. Nat. Genet. 51, 12–18 (2019).

4. Sharifi-Noghabi, H., Zolotareva, O., Collins, C. C. & Ester, M. MOLI: multi-omics late integration with deep neural networks for drug response prediction. Bioinformatics 35, i501–i509 (2019).

5. [1811.06802] PaccMann: Prediction of anticancer compound sensitivity with multi-modal attention-based neural networks. https://arxiv.org/abs/1811.06802.

6. Chiu, Y.-C. et al. Predicting drug response of tumors from integrated genomic profiles by deep neural networks. BMC Med. Genomics 12, 18 (2019).

7. Preuer, K. et al. DeepSynergy: predicting anti-cancer drug synergy with Deep Learning. Bioinformatics 34, 1538–1546 (2017).

8. Sharifi-Noghabi, H., Zolotareva, O., Collins, C. C. & Ester, M. MOLI: multi-omics late integration with deep neural networks for drug response prediction. Bioinformatics 35, i501–i509 (2019).

9. Yang, W. et al. Genomics of Drug Sensitivity in Cancer (GDSC): a resource for therapeutic biomarker discovery in cancer cells. Nucleic Acids Res. 41, D955–D961 (2012).

10. Liu, D. et al. AutoGenome: An AutoML Tool for Genomic Research. bioRxiv 842526 (2019) doi:10.1101/842526.

11. Abadi, M. et al. TensorFlow: A System for Large-Scale Machine Learning. in 12th USENIX Symposium on Operating Systems Design and Implementation (OSDI 16) 265–283 (USENIX Association, 2016).

12. Paszke, A. et al. PyTorch: An Imperative Style, High-Performance Deep Learning Library. in Advances in Neural Information Processing Systems 32 (eds. Wallach, H. et al.) 8026–8037 (Curran Associates, Inc., 2019).

13. Lundberg, S. M. & Lee, S.-I. A Unified Approach to Interpreting Model Predictions. in Advances in Neural Information Processing Systems 30 (eds. Guyon, I. et al.) 4765–4774 (Curran Associates, Inc., 2017).

14. Ashburn, T. T. & Thor, K. B. Drug repositioning: identifying and developing new uses for existing drugs. Nat. Rev. Drug Discov. 3, 673–683 (2004).

15. Wu, Z., Wang, Y. & Chen, L. Network-based drug repositioning. Mol. Biosyst. 9, 1268–1281 (2013).

16. Xu, C. et al. Accurate Drug Repositioning through Non-tissue-Specific Core Signatures from Cancer Transcriptomes. Cell Rep. 25, 523–535.e5 (2018).

17. Langtry, H. D. & Markham, A. Sildenafil. Drugs 57, 967–989 (1999).

18. Yang, W. et al. Genomics of Drug Sensitivity in Cancer (GDSC): a resource for therapeutic biomarker discovery in cancer cells. Nucleic Acids Res. 41, D955–D961 (2013).

19. Chang, Y. et al. Cancer Drug Response Profile scan (CDRscan): A Deep Learning Model That Predicts Drug Effectiveness from Cancer Genomic Signature. Sci. Rep. 8, 8857 (2018).

20. Gao, H. et al. High-throughput screening using patient-derived tumor xenografts to predict clinical trial drug response. Nat. Med. 21, 1318–1325 (2015).

21. McFarland, J. M. et al. Improved estimation of cancer dependencies from large-scale RNAi screens using model-based normalization and data integration. Nat. Commun. 9, 1–13 (2018).

22. Behan, F. M. et al. Prioritization of cancer therapeutic targets using CRISPR–Cas9 screens. Nature 568, 511–516 (2019).

23. Lu, Y.-F., Goldstein, D. B., Angrist, M. & Cavalleri, G. Personalized Medicine and Human Genetic Diversity. Cold Spring Harb. Perspect. Med. 4, (2014).

24. Wirapati, P. et al. Meta-analysis of gene expression profiles in breast cancer: toward a unified understanding of breast cancer subtyping and prognosis signatures. Breast Cancer Res. 10, R65 (2008).

25. Hudson (Chairperson), T. J. et al. International network of cancer genome projects. Nature 464, 993–998 (2010).

26. Bastien, R. R. et al. PAM50 Breast Cancer Subtyping by RT-qPCR and Concordance with Standard Clinical Molecular Markers. BMC Med. Genomics 5, 44 (2012).

27. Rhee, S., Seo, S. & Kim, S. Hybrid Approach of Relation Network and Localized Graph Convolutional Filtering for Breast Cancer Subtype Classification. ArXiv171105859 Cs (2017).

28. Tao, M. et al. Classifying Breast Cancer Subtypes Using Multiple Kernel Learning Based on Omics Data. Genes 10, 200 (2019).

29. Jadaliha, M. et al. A natural antisense lncRNA controls breast cancer progression by promoting tumor suppressor gene mRNA stability. PLOS Genet. 14, e1007802 (2018).

30. Hon, J. D. C. et al. Breast cancer molecular subtypes: from TNBC to QNBC. Am. J. Cancer Res. 6, 1864–1872 (2016).

31. Bogachek, M. V. et al. Sumoylation Pathway Is Required to Maintain the Basal Breast Cancer Subtype. Cancer Cell 25, 748–761 (2014).

32. Hua, H., Zhang, H., Kong, Q. & Jiang, Y. Mechanisms for estrogen receptor expression in human cancer. Exp. Hematol. Oncol. 7, 24–24 (2018).

33. Giacinti, L., Claudio, P. P., Lopez, M. & Giordano, A. Epigenetic Information and Estrogen Receptor Alpha Expression in Breast Cancer. The Oncologist 11, 1–8 (2006).

34. Wang, G., Hao, J., Ma, J. & Jiang, H. A comparative assessment of ensemble learning for credit scoring. Expert Syst. Appl. 38, 223–230 (2011).

35. Koboldt, D. C. et al. Comprehensive molecular portraits of human breast tumours. Nature 490, 61–70 (2012).

